# Mining the forest: uncovering biological mechanisms by interpreting Random Forests

**DOI:** 10.1101/217695

**Authors:** Julian de Ruiter, Theo Knijnenburg, Jeroen de Ridder

## Abstract

Biological datasets are large and complex. Machine learning models are therefore essential to capture relationships in the data. Unfortunately, the inferred complex models are often difficult to understand and interpretation is limited to a list of features ranked on their importance in the model.

We propose a computational approach, called Foresight, that enables interpretation of the patterns uncovered by Random Forest models trained on biological datasets. Foresight exploits the correlation structure in the data to uncover relevant groups of features and the interactions between them. This facilitates interpretation of the computational model and can provide more detailed insight in the underlying biological relationships than simply ranking features. We demonstrate Foresight on both an artificial dataset and a large gene expression dataset of breast cancer patients. Using the latter dataset we show that our approach retrieves biologically relevant features and provides a rich description of the interactions and correlation structure between these features.

## 1 Introduction

The advent of high-throughput genome-wide measurement techniques has allowed researchers to investigate the interaction between many biological components in an organism as well as their effect on phenotype. Such studies frequently depend on computational modeling as the sheer amount of variables and data is beyond what the human intellect can appreciate. Many successful computational models have been developed allowing for such feats as cancer sub-typing [14], survival prediction [12] and personalized medicine recommendation [18] - all based on large and heterogeneous datasets. On the downside, the predictive power of these models is often in sharp contrast with the domain expert’s ability to understand them. This is because most inference methods result in ‘black-box models’ that lack a good biological interpretation. In most cases a simple list of features, ranked on their importance in the model is the only output that is accessible to the domain expert. This limited interpretability impedes the formulation of novel hypotheses or follow-up experiments.

One strategy to address this challenge is to limit the complexity of the model. For instance, one could - a priori - severely reduce the number of features in the model by means of feature selection or cluster averaging to ensure the final predictor remains interpretable. Alternatively, model complexity can be reduced by restricting the model class to linear models, possibly combined with regularization such as the lasso [26] and group lasso [30], thereby allowing some interpretation by inspection of the regression coefficients. However, evidence is mounting that complex non-linear models, such as Random Forest models, are much better at fitting to the data [5, 19]. Such models are able to adapt to structure in the data that is not captured using linear models or by models in a reduced feature space. In this work we therefore take an alternative approach, that is, instead of limiting the complexity of the model beforehand we aim to inspect the structure of a complex non-linear model to enable interpretation of the underlying biology captured by the model. Our approach, called Foresight, is tailored to interpretation of a Random Forest (RF) model trained on a large collection of features and summarizes interactions between informative features in a network representation. Importantly, Foresight is not a feature selection method, but a post-hoc analysis, which derives relevant features and their interactions from trained RFs.

The Random Forest is an ensemble learning method [3] which has gained much popularity in the analysis of biological datasets due to its high prediction accuracy in noisy, high dimensional and under-sampled classification tasks, typical for biological problems [27]. It is also a clear example of a black-box model, as an RF consists of thousands of individual decision trees. As a result, RFs are difficult to interpret. The most widely used method to interpret RFs is to rank individual features by the RF’s variable importance score (see the Extended Methods and materials in the Supplemental Information for more details) and to select the top features [10]. This has been successfully exploited in the identification of gene signatures for predicting drug-sensitivity [20] and detection of prognostic markers for recurrence of hepatocellular carcinoma [29].

A significant challenge for the post-hoc interpretation of RFs and other classification models is the complex patterns of correlation between features [10]. The reason for this is that correlated features are effectively interchangeable within the model. This causes dilution of importance scores within a correlated group of features. As a result, models are likely to attribute high importance to only one or a few features from a correlated group of features. This can potentially lead to ambiguous interpretation because small differences in the underlying dataset give rise to entirely different sets of important features. This is a well-known impediment for the biological interpretation of signature genes for breast cancer outcome prediction as different studies report different gene sets with only minimal overlap [15, 8].

To deal with this challenge, Foresight takes the correlation structure between features into account by diffusing importance scores between correlated features and correlated interacting feature pairs. This diffusion process has the effect of boosting the importance scores of correlated predictive features, thus counter-acting the dilution of importance scores within correlated groups. Furthermore, by boosting the scores of the entire group, this approach is much more likely to select the majority of the features in a correlated feature group. This is of significant advantage in downstream analyses of biological datasets as this allows us to identify biological pathways and/or functions that are overrepresented in high-ranking genes.

We demonstrate the utility of our approach by applying it both to an artificial dataset and to the METABRIC dataset, one of the largest gene expression datasets of breast cancer samples to date [6]. In the latter dataset the objective is to differentiate between two prominent types of invasive breast cancer: invasive ductal carcinoma (IDC) and invasive lobular carcinoma (ILC). It is important to note that, as ILCs can be readily distinguished from IDCs using histopathology, the goal of this classification is not to predict cancer subtypes for new patients. Instead, we are interested in delineating the differences between the subtypes at the molecular level. Understanding the molecular mechanisms that determine the difference between ILC and IDC could facilitate a tailored treatment that targets each of the distinct pathological subtypes specifically, ultimately leading to benefits for the patient. In this work, we show how interpretation of this dataset using Foresight can help to reach these important goals.

## 2 Approach

Foresight’s approach consists of four steps, which are graphically depicted in Figure 1 and are explained in more detail in the Methods. Briefly, in the first step, a trained RF model is used to calculate support scores for both individual features and pairs of features (Figure 1A). For single features, the support score is based on the number of times an individual feature occurs in the forest. For pairs of features, this score is based on the number of times two features co-occur in a tree of the forest. Second, support scores are diffused across correlated features and correlated feature pairs using a *k*-nearest-neighbor diffusion approach (Figure 1B). This counteracts the dilution of support scores across many correlated features. Third, the individual features and feature pairs are ranked based on their diffused support scores (Figure 1C) and the highest ranking ones are selected. In the final step, the selected features are visualized in a network representation (Figure 1D). This representation facilitates interpretation by highlighting both feature co-occurrence, which indicate a synergistic relation between the corresponding biological features, and feature correlation, which points to a similar biological role for these features.

**Fig. 1.**
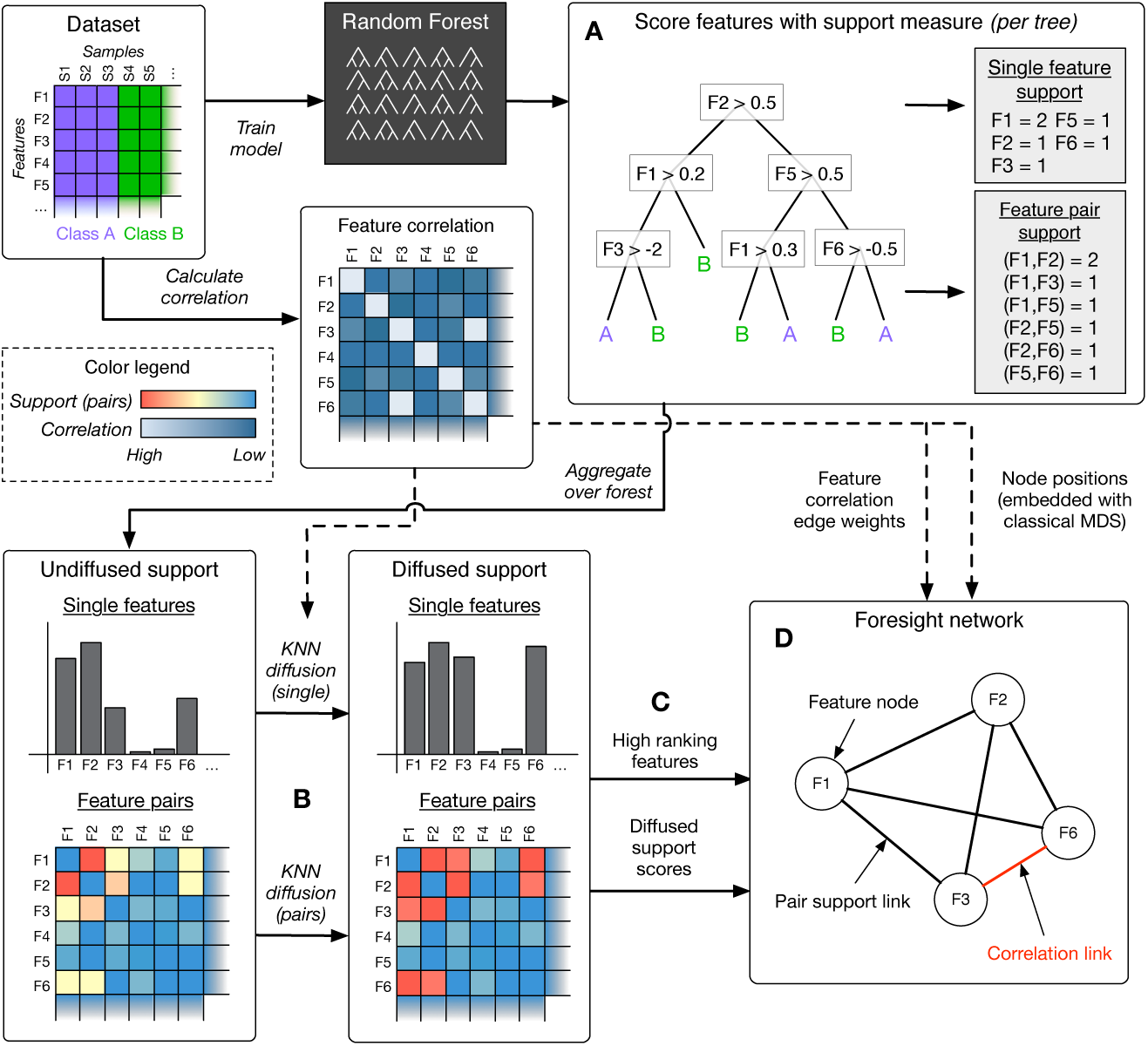
Overview of Foresight’s approach. (A) Features and feature pairs are scored using the support score. (B) The scores are adapted using diffusion to account for correlation between features. (C) The feature (pairs) are ranked based on the diffused support scores. (D) The highest-ranking features are visualized in a network representation. In this example features F1, F2, F3 and F6 are equally predictive and only features F3 and F6 are correlated.

## 3 Methods

### 3.1 Scoring of features and feature pairs

Individual features and feature pairs are scored using the support score. For single features this score is defined as the number of times the feature is used at a decision node (non-leaf node) in trees of the RF. Thus, the single feature support score (S) of a feature *f* is defined as:

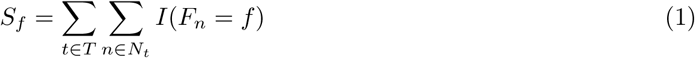

in which *T* represents the collection of all decision trees *t* in the forest, *N*_*t*_ the collection of all non-leaf nodes *n* of the decision tree *t* and *F*_*n*_ the feature used in node *n*. A high value for the score indicates that the feature is highly predictive, as it is frequently used in the predictions of the individual trees in the forest. A low support score means that the feature is rarely used in the prediction process and therefore of little predictive value.

Similarly, the feature pair support score (PS) is defined as the number of times that two features co-occur at decision nodes along a single path from the root of a tree to one of its leaves. As such, the support of a feature pair (*f, g*) is defined as:

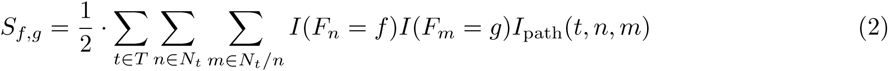

with *I*_path_(*t, n, m*) = 1 if nodes *n* and *m* occur in a single path from the root of tree *t* to one of its leaves and 0 otherwise.

As an example, Figure 1 states both types of support score for a single decision tree. Note that features F5 and F6 are counted as a co-occurring feature pair as they occur along the same path from the root of the tree to its right-most leaf. However, features F1 and F6 do not form a feature pair, as they do not co-occur in any path from root to leaf. The feature pair (F1, F2) occurs twice in the tree, once in the left half of the tree and once in the right, and is therefore assigned a support score of 2.

### 3.2 Diffusion of scores within correlated neighborhoods

Highly correlated features are effectively interchangeable for classification. As a result, the support scores within a group of highly correlated features are diluted among the members of the group, thus lowering the scores of the individual features. In Foresight this effect is counteracted by aggregating contributions from features within the correlation neighborhood of each feature through a diffusion step. This step effectively boosts the support scores of correlated features to better reflect their true predictive value. We use a *k*-nearest-neighbor approach such that the same number of features (*k*) contributes support scores to each feature. These contributions are weighted based on correlation. The resulting diffused support score (DS) is defined as:

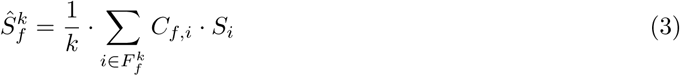

with *C*_*f,i*_ denoting the absolute Pearson correlation weight between features *f* and *i* and the set 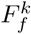 representing set of *k* features closest to feature *f* in terms of their correlation. Note feature *f* is itself also present in this set.

Support scores for feature pairs are diffused in a similar manner. Here, the correlation neighborhood of a feature pair (*f, g*) is defined based on the product of the individual feature correlations to features *f* and *g*. Specifically, the diffused pair support (DPS) is defined as:

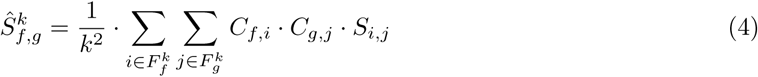

Again, *f* and *g* are included in *F*_*f*_ and *F*_*g*_, respectively. This ensures that the support score between *f* and *g* will contribute most towards 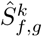, because *C*_*f,f*_ and *C*_*g,g*_ are 1.

Note that we defined the extent of the diffusion of the score of a feature based on its *k*-nearest-neighbors instead of based on the correlation scores of all other features. Thus, the diffusion neighborhood is of equal size for features and feature pairs with many correlates and those with just a few. Moreover, in this way we do not have to rely on a strict clustering of features.

The choice of *k* involves a trade-off between limited diffusion (low values of *k*), which potentially masks relevant features and feature pairs, or high diffusion (i.e. high values of *k*), which may include contribution of uncorrelated neighbors and may bias for larger groups of correlated features. In this work we use *k* = 15, but show that the results are relatively insensitive to its setting.

### 3.3 Ranking of features and feature pairs

The (diffused) support scores are used to rank features and feature pairs based on their importance in the RF. The resulting ranking is used to identify the top features and feature pairs, which are selected for visualization in the final step of the approach.

In the following, we will compare four different scores that are based on the (diffused) support scores. Table 1 provides an overview of these scores along with the variable importance score (VI). The VI score is included for later comparison with the support scores, since it is the default score used to rank features using RFs. It is based on the mean decrease in Gini impurity, and is calculated as part of the training procedure of the RF (see Methods and materials for more details). Abbreviations for each of the scores are listed in Table 1, which we will use in the remainder of the paper.

**Table 1.** An overview of the scores used to rank single features and feature pairs.

| Name | Abbr. | Type | Equation | Ranking single | Ranking pairs |
| --- | --- | --- | --- | --- | --- |
| Support | S | Single | 1 | Used directly | Minimum score of members in the pair |
| Diffused support | DS | Single | 3 |  |  |
| Variable Importance | VI | Single | S1 |  |  |
| Pair support | PS | Pairwise | 2 | Flatten pair list, rank by order in flattened list | Used directly |
| Diffused pair support | DPS | Pairwise | 4 |  |  |

To allow for a comparison between the single feature scores (S, DS and VI) and the feature pair scores (PS and DPS), we convert the single scores to pair scores by calculating the score of each individual feature in a pair and taking the minimum of these two scores as the pair score. For example, for the VI we obtain a feature pair score as: VI_*f,g*_ = min(VI_*f*_, VI_*g*_). Reciprocally, single feature rankings are obtained from pair scores by first using the pair score to rank feature pairs and subsequently ranking the single features by the order in which they appear in the ranked pair list.

### 3.4 Visualization of feature correlation and interaction for interpretation

The final step of Foresight’s approach is to select the top ranking features and feature pairs, and to visualize these in a network representation that highlights interaction and correlation between features. Pairs of features are considered to interact if they are frequently selected together by the RF. This means that the specific combination of these two features is beneficial for accurate classification. In a cancer-related dataset, such a pair of features may represent two separate biological processes that must be perturbed simultaneously for the development of the cancer. On the other hand, correlation between features points to a similar role for these features. For example, two highly correlated features may be expected to function in a single signaling pathway or protein complex and therefore play a similar role in tumorigenesis. By visualizing both these relations in a single network representation, we allow the user to identify groups of highly correlated features and, importantly, simultaneously identify strong interactions between these groups.

Features are represented as nodes in the network and may be connected by two different types of edges: correlation edges and interaction edges. Two feature nodes are only connected by a (red) correlation edge if the correlation between their features exceeds the preset minimum correlation threshold. Similarly, two nodes are only connected by a (black) interaction edge if the support value of their interaction pair exceeds the predefined minimum support threshold. Furthermore, the transparency of the edges is used to visualize the relative magnitude of the co-occurrence or correlation, with higher scores corresponding to more opaque edges.

Nodes in the network are positioned based on their correlation, such that highly correlated features manifest as groups and uncorrelated features as distinct entities. This approach adds considerable clarity to the resulting network by allowing edges in the network to be analyzed at the level of grouped features instead of single features. Foresight determines the positions of nodes in the network using classical multidimensional scaling to reduce the matrix of absolute feature correlation distances to a two dimensional space.

## 4 Results

### 4.1 Identification of predictive features in artificial data

**Construction of the artificial dataset** To test the performance of our approach, we created a realistic artificial dataset, with a known ground truth, based on a colon cancer gene expression dataset [17]. The top 5000 most variable genes in the gene expression data were selected as features for the artificial dataset. The features were clustered using the correlation distance measure to identify clusters of highly correlated features. Three of these clusters, containing 7, 17 and 14 correlated features respectively, were selected to define the ground truth of the dataset. These clusters will hence be referred to as *X*_1_, *X*_2_ and *X*_3_. Using these clusters, the class label of each sample was defined as:

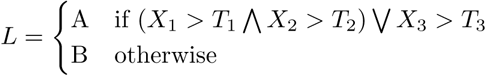

in which *X*_*i*_ > *T*_*i*_ is only true if all features in cluster *X*_*i*_ exceed the predefined thresholds specified in *T*_*i*_. Note that *T*_*i*_ contains a threshold for each feature in *X*_*i*_, which were determined emperically to balance the size of the classes.

This construction ensures that the dataset contains many clusters of correlated features, of which only the three selected clusters are truly predictive. As such, the dataset mimicks a potential biological situation in which three groups of co-expressed genes together result in a specific phenotype. Furthermore, as the features are based on actual gene expression data, both the size and composition of the clusters in the dataset reflect that of an actual biological dataset.

#### Evaluation of scores for ranking single features

We assessed the performance of the (diffused) support scores and the VI in the selection of individual features by ranking the features in the artificial dataset on each of the scores and comparing the resulting rankings against the known ground truth (consisting of the features in clusters *X*_1_, *X*_2_ and *X*_3_). The rankings were scored against the known ground truth of the dataset by calculating the receiver operating characteristic (ROC) and the corresponding area under the curve (AUC). To emphasize the importance of identifying ground truth features early in the ranked list, AUCs were only calculated for FPR *<* 0.1. This AUC is subsequently multiplied by to scale the score between 0 and 1. We refer to this score as the AUC01.

The robustness to noise of each of the scores was evaluated by repeating this procedure with different levels of noise in the dataset. The amount of noise in the dataset was defined by a parameter *α*, with *α* = 0 corresponding to a situation without noise and *α* = 1 to a dataset only consisting of noise (see the Extended Materials and methods in the supplement for more details). The variability in the calculated AUCs was determined by repeating the analysis for 10 instances of the artificial dataset at each noise level, in which only the stochastic noise differs between instances. Note that the noise level also reflects the difficulty of classification for the RF model, resulting in the trained models having a mean out-of-bag (OOB) prediction error of approximately 0.14 for *α* = 0 and OOB errors of 0.16, 0.18 and 0.44 for *α* = 0.5, *α* = 0.75 and *α* = 1, respectively.

The resulting AUC curves for single features are depicted in Figure 2A, with the ROC curves for a single instance with *α* = 0.5 shown in Figure 2B. From the AUCs it is evident that the DPS outperforms the other measures for *ff <* 0.75, indicating that it is indeed beneficial to include pair information in the ranking of individual features. We also find that the DS outperforms the S and the VI marginally for low noise levels with its advantage increasing at higher levels of noise. From the ROC curve (shown in Figure 2B) we see that the advantage of the DS is mainly attributable due to its overall higher true positive rate (TPR), as for lower false positive rates (FPR) the score lags behind the S and the VI. This indicates that the main benefit of the single feature diffusion comes from ensuring that all features from the three clusters are selected, whereas the S and VI measures fail to select some of the predictive features.

**Fig. 2.**
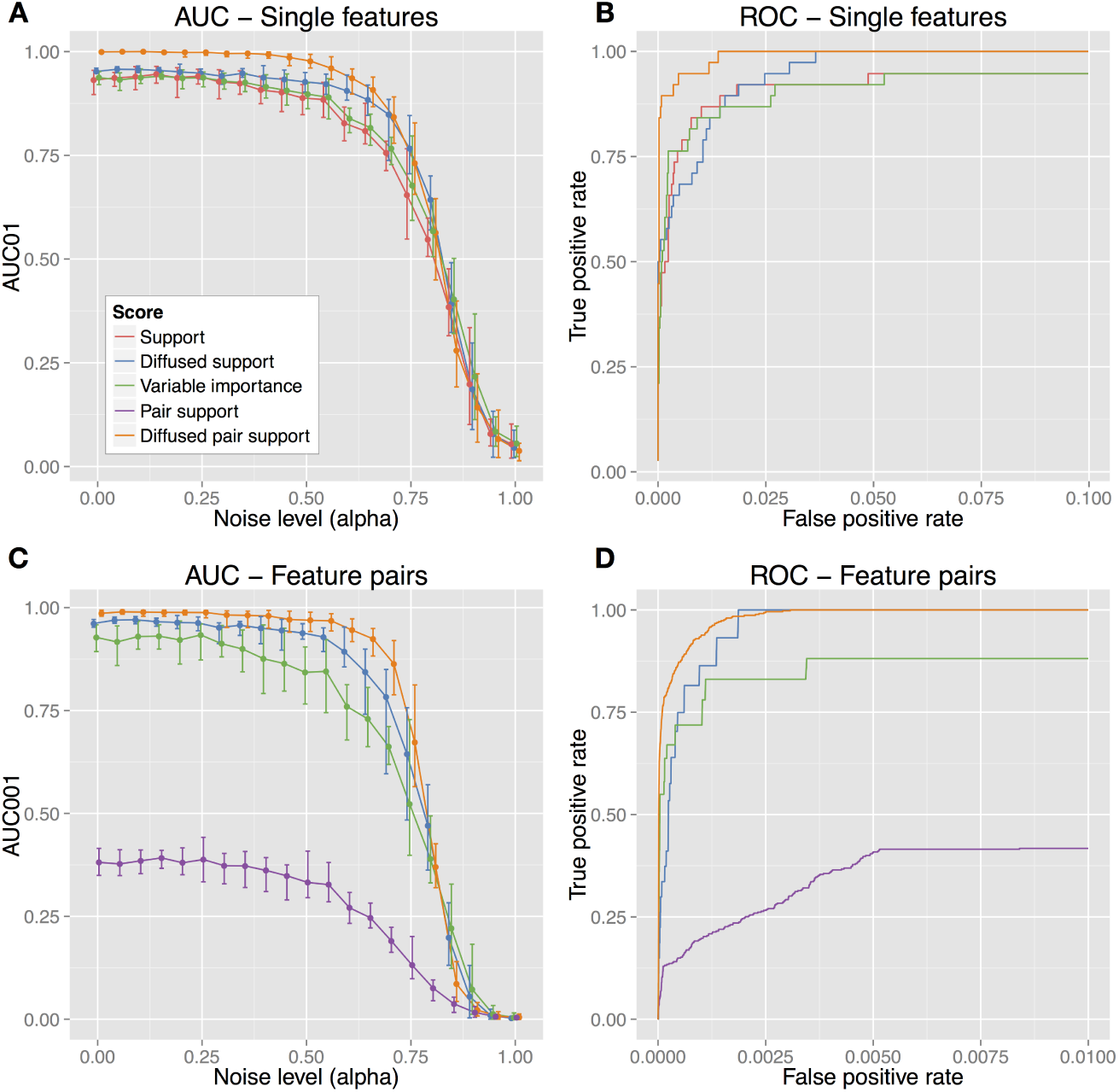
AUC and ROC curves for the selection of single features and feature pairs in the artificial dataset. Figures A and B respectively depict the AUC and ROC curves for the selection of individual features. Figures C and D shown similar curves for the selection of feature pairs. The AUC curves demonstrate the performance of the various scores across different noise levels, specified by noise parameter *α*. Both ROC curves were created with *α*= 0.5 to illustrate score performance at an intermediate noise level.

#### Evaluation of scores in the ranking of feature pairs

The same approach was used to assess the performance of the scores in the ranking of feature pairs. For feature pairs we defined the ground truth >of the artificial dataset as all interactions between the three ground truth clusters (i.e. all combinations of features *X*_1_-*X*_2_, *X*_1_-*X*_3_ and *X*_2_-*X*_3_), as these pairs constitute all interactions between the three predictive clusters. As the amount of feature pairs is much larger than the amount of individual features, the rankings of feature pairs are scored using the AUC001, which is similar to the AUC01 but limits it calculation of the AUC to FPR *<* 0.01 and is multiplied by 100 to scale the score between 0 and 1.

From the feature pair AUCs (Figure 2C) we see that the DPS notably outperforms the other measures up to a noise level of 0.75, which is a similar result to that of the single feature case. Though the VI performs reasonably, it is outperformed by the diffused scores. This is due to its failure to select all predictive feature pairs, as is evident from the ROC curve (Figure 2D). Interestingly, the pair support measure performs poorly, showing that the diffusion step is crucial for the performance of the diffused pair support.

Overall these results demonstrate that the DPS outperforms the S, PS, DS and VI scores in both the selection of single features and feature pairs. This shows that both the diffusion and the inclusion of pair information contributes considerably to accurate feature selection in this artificial dataset. This advantage is only lost at high noise levels (*α >* 0.75), likely due to the loss of correlation and/or loss of informative features at these high noise levels. Furthermore, additional experiments have shown that this result is robust across different sizes of the RF (Suppl. Figure S1) and different values of the KNN diffusion parameter *k* (Suppl. Figure S2).

#### Recapitulation of dataset structure using the network visualization

To determine if our network visualization is able to recapitulate the structure underlying the artificial dataset, we used the DPS to select and visualize the strongest feature pairs in the dataset. Figure 3 shows the result after minimal manual manipulation (the raw Cytoscape visualization is given in Figure S6). From the visualization the three ground truth clusters *X*_1_, *X*_2_ and *X*_3_ can easily be identified as the three distinct groups of features that are highly connected by correlation edges within their group. Furthermore, the large amount of black edges between the three groups clearly shows a high degree of co-occurrence between the features of the respective groups, reflecting the importance of combining features from the three groups for accurate prediction. Altogether this shows that this approach can clearly identify the feature structure embedded in the artificial dataset.

**Fig. 3.**
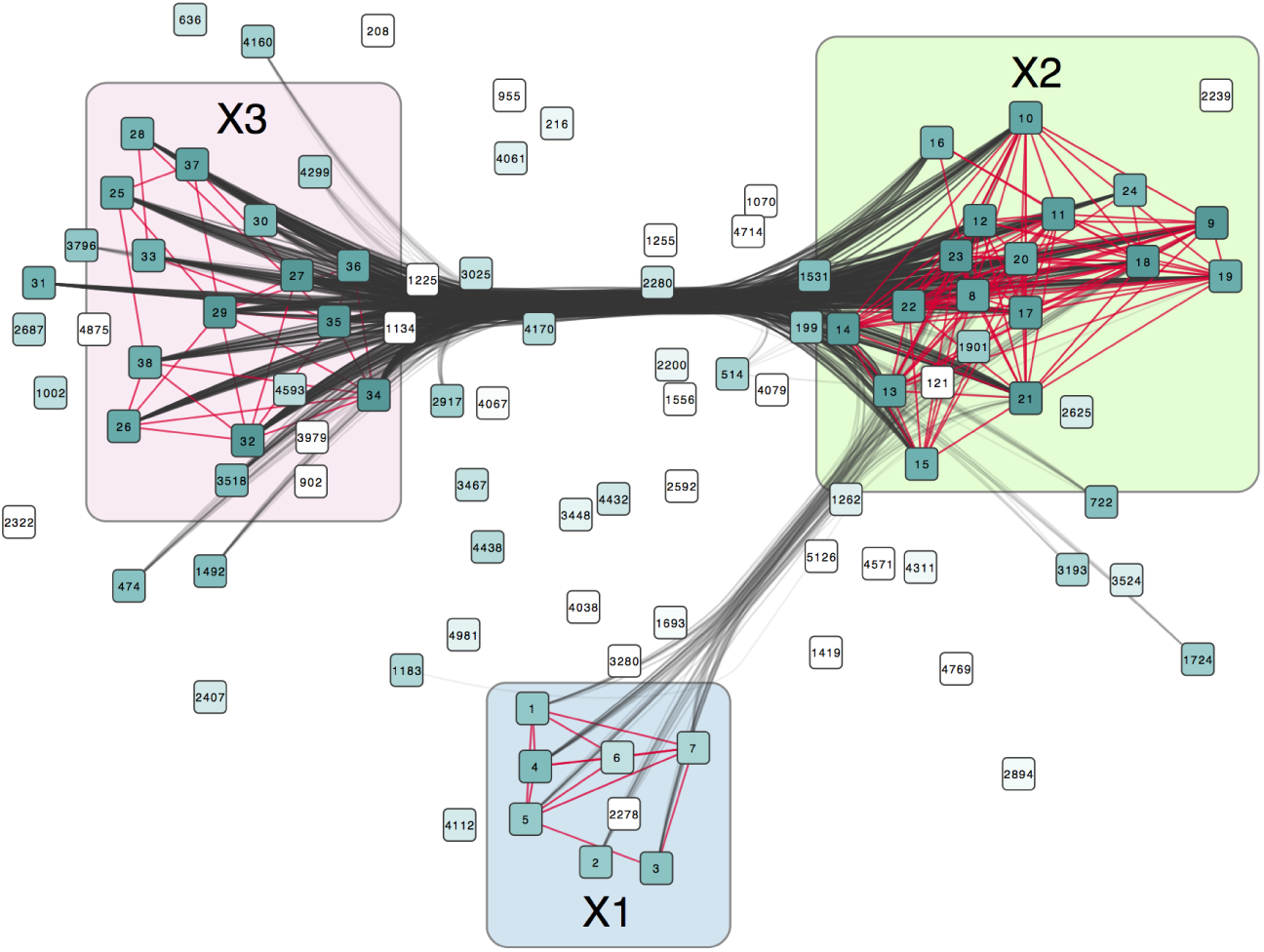
Foresights network representation of the top 100 features (ranked by the DPS) in an instance of the artificial dataset with *α*= 0.2. Nodes in the network correspond to the individual features and are colored based on the rank of their feature, with a darker color corresponding to a higher rank. A red correlation edge connects two nodes if their correlation exceeds 0.8. Similarly, nodes are connected by a black feature pair edge if their pair support exceeds 75. These two thresholds were chosen to balance the information content and the sparsity (and thus the interpretability) of the network. The transparency of both edge types indicates their relative strength in the network. The locations of the predictive features in clusters *X*_1_, *X*_2_ and *X*_3_ are indicated by their respective colored regions, showing that this network representation provides a clear view of both the correlation structure within each cluster and degree of support between two clusters.

### 4.2 Interpretation of features that discriminate between ILC and IDCs in METABRIC

After establishing the usefulness of Foresight on artificial data, we set out to determine if our approach could facilitate the interpretation in an actual dataset. To this end, we applied Foresight to a RF trained to discriminate ILC and IDC breast cancer subtypes based on gene expression. We evaluated the ranked features and feature pairs by determining enrichment in gene sets. Gene set enrichment analyses, for example GSEA [16, 24] and DAVID [11], have become a standard approach for high level functional interpretation of gene lists. We used the number of enriched gene sets as a measure of interpretability. Furthermore, we evaluated the results by overlaying them with protein-protein interaction networks, and through inspection of the network visualization.

The ILC vs. IDC gene expression dataset is a subset of the METABRIC dataset. We randomly selected a total of 294 tumors, which were spread evenly over both classes to avoid biasing the RF towards either of the classes. Non-informative features were removed by selecting the top 5000 most variable features as input for the analysis. This dataset was used to train a RF of 2000 trees with a reasonably high prediction accuracy, as indicated by the model’s OOB error of 0.21.

#### Identification of relevant enriched gene sets with GSEA

The diffused pair support score (DPS, Eq. 4) was used to rank the features in the dataset, after which we applied GSEA to determine if the ranking was enriched for breast-cancer or, more specifically, IDC/ILC related gene sets. The same GSEA analysis was also performed on a gene list ranked by the VI score for comparison with the VI-based approach.

The analysis revealed that the DPS finds substantially more enriched gene sets than the VI, identifying 123 enriched gene sets with FDR *<* 0.1 versus only 2 found by the VI. We further compared the top ten gene sets of both approaches (ranked by the FDR) and used boxplots to visualize the distribution of the ranks of the genes in each gene set (Figure 4). Overall, the ranks of the genes in the DPS top gene sets (Figure 4A) are considerably better than those of the sets identified by the VI (Figure 4B). Moreover, within the selected gene sets the rankings of the DPS are better than those of the VI. This explains, to a large extent, the difference in the enrichment between the two rankings and demonstrates that the DPS does indeed enrich for functionally coherent sets of genes. This analysis clearly shows the effect of taking into account both the correlation structure and the analysis of pairs of features.

**Fig. 4.**
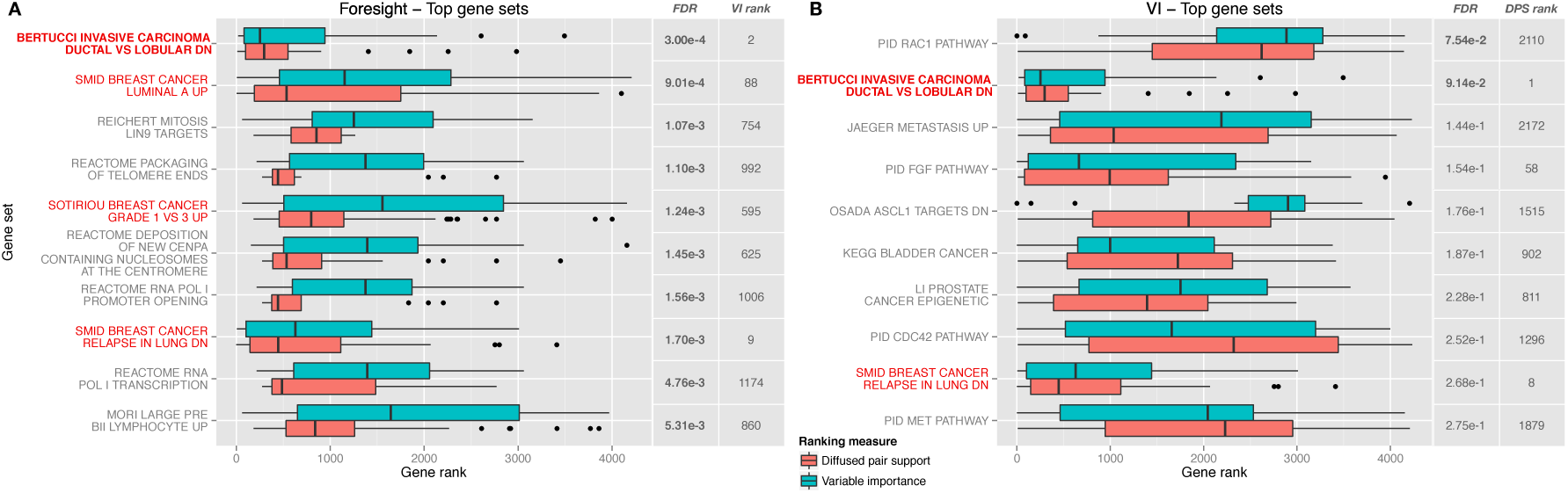
Top 10 gene sets identified by Foresights DPS (Figure A) and the VI (Figure B) Gene sets clearly related to breast cancer are highlighted in red. For each gene set we show a box plot of the ranks of the genes in the gene set to illustrate how the genes are ranked by the DPS and VI respectively. The FDR of each gene set and its rank in the other approach is shown for comparison. The FDRs of significantly enriched gene sets (*FDR <* 0.1) are indicated in bold. From the FDRs and the overall gene ranks within the gene sets it is clear that Foresight identifies more enriched gene sets than the VI.

To determine if Foresight’s diffusion specifically enriches for gene sets that are relevant to the distinction between ILCs and IDCs, we examined the selected top gene sets for their relevance to breast cancer and/or the discrimination between ILCs and IDCs. Of the top ten Foresight gene sets, four can be directly related to breast cancer. The highest ranking gene set (Bertucci) corresponds to a gene expression signature that was previously determined to distinguish ILCs and IDCs using microarray gene expression data [1]. The idenification of the Smid luminal A gene set (rank 2) is in line with the observation that ILCs are generally luminal A breast cancers, whilst IDCs are typically more diverse [13]. Similarly, ILCs tend to have a lower grade than IDCs [21], explaining the enrichment of the Sotirou grade gene set (ranked no. 5). The bias for lower grades and the luminal A subtype were indeed confirmed from the clinical data of the samples (Supplementary Tables S1 and S2). Additionally, the presence of the Smid relapse gene set is likely due to the differences in metastatic spread between ILCs and IDCs, as IDCs are more often found to form lung metastases than ILCs [9].

The VI approach also identifies the Bertucci and Smid relapse gene sets as top hits that are directly related to breast cancer, however only the Bertucci set is statistically significant with FDR *<* 0.1 (FDR = 9.14·10^-2^, compared to 3.00·10^-4^ in the Foresight ranking). The relevance of other genes sets in the top 10 is unclear. For example, though the RAC1 pathway has been associated with migratory and invasive behavior [7], its enrichment is weak and stems solely from the high ranks of CDH1 and MAP3K1. Additionally, although the FGF, CDC42 and MET pathway gene sets are attributed to invasive behavior and metastasis in the literature [25, 23, 2, 22], their role specific to IDCs and ILCs is unclear and their set enrichment and the overall gene ranks are underwhelming.

In summary, these results support the idea that the DPS score in Foresight enables a better interpretation of the METABRIC ILC vs. IDC dataset than the traditional VI. Specifically, ranking of the genes based on DPS leads to more significant gene sets, many of which are directly relevant to the biological problem under investigation.

#### Visualization and interpretation of structure between predictive features

To gain additional insight into the correlation structure and interaction between features in the METABRIC dataset, we selected the top 100 features from the ranked list and visualized these features using Foresight’s network representation. The resulting network (shown in Figure 5, the raw Cytoscape visualization is provided in Figure S7) contains 85 unique genes and can clearly be divided into two groups of correlated features and a small group of features without high-ranking correlates. Of the 85 genes in Figure 5, we identified five genes (CDH1, SPRY1, CCL14, SPP1 and THBS4) that have previously been identified to distinguish ILCs and IDCs by Turashvili et al. [28]. Another six genes (MFAP4, SPARCL1, ELN, ALDH1A1, DPT and CD34) are included in the Bertucci ILC vs. IDC gene signature [1] that was enriched in our GSEA analysis. Additionally, genes such as GJB2, COL11A1, MMP11, PLAUR and PLAU have been associated with the development of IDCs[4]. Taken together, these results show that many genes in the network are known to play a role in the development of ILCs and IDCs.

**Fig. 5.**
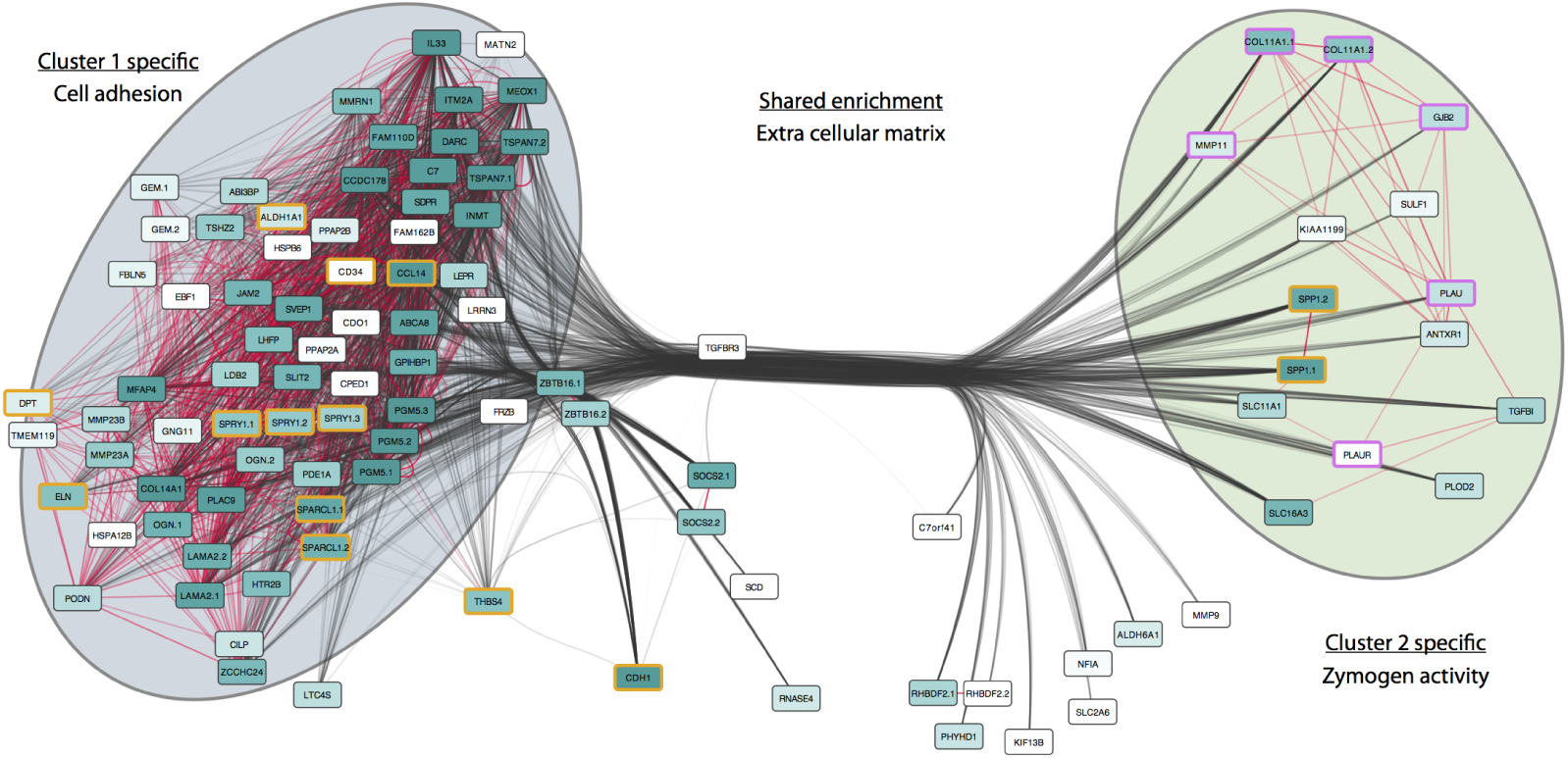
Network visualization of the top 100 features in the METABRIC ILC vs. IDC dataset (as ranked by the diffused pair support) Nodes are labeled by the gene symbol that corresponds to the Illumina probe ID of their feature, with numbered suffixes added for genes with multiple probes. Node colors correspond to the relative rank of their features, ranging from dark green (high) to white (low). Highly correlated features are connected by red links and highly scoring feature pairs are indicated by black links, with the transparency of the links indicating their relative strength. Genes indicated in orange correspond to genes that are known to distinguish ILCs and IDCs, those colored in purple are known to play a role in the development of IDCs.

The topology of the network also reveals clear substructure between the selected features. There appear to be two main clusters, which are not only different in terms of their gene expression (Supplementary Figures S3 and S4) but also heavily connected by support edges. This means that features from both clusters frequently co-occurred in the RF, indicating that combinations of features from these clusters are important in discriminating between IDCs and ILCs. The high functional enrichment observed when ranking genes based on the DSP prompted us to investigate whether gene pairs that were highly ranked by DSP were indeed functionally related. Using a shortest path distance approach between gene pairs in protein-protein interaction networks, we determined that, indeed, gene pairs selected by Foresight are connected by shorter paths than pairs selected by other approaches such as the VI (Suppl. Figures S8 and S9).

Interestingly, DAVID [11] reveals cluster-specfic functional enrichment (summarized in Figure 5, detailed results in Supplementary Files S1 and S2). This observation directly points to the possible interaction between biological functions important in the context of ILCs and IDCs, which warrants further investigation by domain experts.

## 5 Discussion

The utility of complex computational models in biology is not limited to their predictive performance or data fit, but also the extent to which they can help domain experts generate new hypotheses or experiments. This seems somewhat paradoxical as these complex models exceed the human intellect and are thus considered black-box models. Therefore, post-processing steps have to be taken to extract useful information from the model. For example, in most models importance scores can be calculated, allowing the identification of the most informative features.

If these features are genes, a common additional post-processing step is gene set enrichment analysis. This allows long lists of genes to be summarized by a small number of enriched gene sets that represent the biological functions underlying the gene list. This greatly helps in providing the domain expert insight into the model and explains the large appeal and wide use of these approaches.

Here, we have introduced a post-hoc analysis for the Random Forest, a classification model that has gained much traction in the field of computational biology due to its robustness and overall high prediction accuracy. Our approach, called Foresight, facilitates the interpretation of RF models by identifying and visualizing the informative features and their interactions. Central to this approach is the use of the correlation structure between the features and the identification of important feature pairs in the RF.

Using the METABRIC ILC vs. IDC dataset, we have shown that this approach allows us to identify more enriched gene sets that are relevant to the underlying biology than the traditional variable importance score. Furthermore, selection and visualization of high scoring feature pairs in a network representation, enabled us to identify correlated groups of features with distinct biological properties. These results exemplify the contribution of Foresight, which is to facilitate domain experts in the interpretation of RFs applied to large datasets.

We observed that exploiting pairs of features outperforms the use of single feature scores as well as the more sophisticated VI score to retrieve the true relationships in the data. While it is computationally expensive to analyze feature pairs, with current computation power this is certainly not prohibitive. In future it may even become straightforward to analyze feature pairs and possibly even feature triplets.

Although our approach is practical and robust, several important considerations must be mentioned. First, a potential drawback of the diffusion approach to boost the scores of correlated features is that it may inadvertently bias feature scores as a function of the size of the correlated group. Our results indicate, however, that important players in ILC vs. IDC differentiation, such as CDH1, were without correlates in the data, yet still discovered by our approach. Nevertheless, it is important to investigate to what extent group size biases diffused scores and if other measures than the Pearson correlation coefficient may be more suitable.

Second, there can be alternative reasons why feature pairs are jointly selected in trees in a RF other than that they contain different information and that their combination is informative for the class labels. For example, it is also possible that they contain similar information and their combination provides robustness for classification, in which case the features will be both highly correlated and co-occurrent. Untangling these types of pairwise relations will provide considerably more insight into the manner in which features are used in the classification model.

Third, the current approach provides a general view of the structure of the RF as scores are aggregated over the entire forest, but does not take structure between samples into account (other than through the class labels). Analyzing the RF on a per-sample basis or for subsets of samples may be valuable as it will allow the distinction of groups of predictive features that explain distinct subsets of samples in the dataset. This would be especially relevant in the analysis of biological datasets, considering the heterogeneous nature of biological samples.

An important avenue for further study is to investigate if other types of black box models are amenable to interpretation as well. We believe that the underlying principle of Foresight, i.e. a co-occurrence analysis whilst taking into account the correlation structure in the data, is readily applicable to other complex models. This will be essential for leveraging the full potential of the vast amounts of data that are currently being generated in the fields of molecular medicine and the life sciences in general.

## Acknowledgement

This study makes use of data generated by the Molecular Taxonomy of Breast Cancer International Consortium. Funding for the METABRIC project was provided by Cancer Research UK and the British Columbia Cancer Agency Branch. J. de Ridder is supported by the Netherlands Organisation for Scientific Research (NWO-Veni: 639.021.233).

## References

1. F Bertucci, B Orsetti, V Nègre, P Finetti, C Rougé, J-C Ahomadegbe, F Bibeau, M-C Mathieu, I Treilleux, J Jacquemier, L Ursule, A Martinec, Q Wang, J Bénard, A Puisieux, D Birnbaum, and C Theillet. Lobular and ductal carcinomas of the breast have distinct genomic and expression profiles. Oncogene, 27(40):5359–5372, September 2008.

2. Kristi Bray, Melissa Gillette, Jeanette Young, Elizabeth Loughran, Melissa Hwang, James Sears, and Tracy Vargo-Gogola. Cdc42 overexpression induces hyperbranching in the developing mammary gland by enhancing cell migration. Breast cancer research : BCR, 15(5):R91, September 2013.

3. Leo Breiman. Random Forests. Machine Learning, 45(1):5–32, 2001.

4. Bàrbara Castellana, Daniel Escuin, Gloria Peiró, Bárbara Garcia-Valdecasas, Tania Vázquez, Cristina Pons, Maitane Pérez-Olabarria, Agustí Barnadas, and Enrique Lerma. ASPN and GJB2 Are Implicated in the Mechanisms of Invasion of Ductal Breast Carcinomas. Journal of Cancer, 3:175–183, 2012.

5. James C Costello, Laura M Heiser, Elisabeth Georgii, Mehmet Gönen, Michael P Menden, Nicholas J Wang, Mukesh Bansal, Petteri Hintsanen, Suleiman A Khan, John-Patrick Mpindi, et al.A community effort to assess and improve drug sensitivity prediction algorithms. Nature Biotechnology, 2014.

6. Christina Curtis, Sohrab P Shah, Suet-Feung Chin, Gulisa Turashvili, Oscar M Rueda, Mark J Dunning, Doug Speed, Andy G Lynch, Shamith Samarajiwa, Yinyin Yuan, Stefan Gräf, Gavin Ha, Gholamreza Haffari, Ali Bashashati, Roslin Russell, Steven McKinney, METABRIC Group, Anita Langerød, Andrew Green, Elena Provenzano, Gordon Wishart, Sarah Pinder, Peter Watson, Florian Markowetz, Leigh Murphy, Ian Ellis, Arnie Purushotham, Anne-Lise Børresen-Dale, James D Brenton, Simon Tavaré, Carlos Caldas, and Samuel Aparicio. The genomic and transcriptomic architecture of 2,000 breast tumours reveals novel subgroups. Nature, 486(7403):346–352, June 2012.

7. J Deplazes, M Fuchs, S Rauser, H Genth, E Lengyel, R Busch, and B Luber. Rac1 and Rho contribute to the migratory and invasive phenotype associated with somatic E-cadherin mutation. Human Molecular Genetics, 18(19):3632–3644, September 2009.

8. Liat Ein-Dor, Itai Kela, Gad Getz, David Givol, and Eytan Domany. Outcome signature genes in breast cancer: is there a unique set? Bioinformatics, 21(2):171–178, January 2005.

9. S Ferlicot, A Vincent-Salomon, J Médioni, P Genin, C Rosty, B Sigal-Zafrani, P Fréneaux, M Jouve, J P Thiery, and X Sastre-Garau. Wide metastatic spreading in infiltrating lobular carcinoma of the breast. European Journal of Cancer, 40(3):336–341, February 2004.

10. R Genuer, J M Poggi, and C Tuleau-Malot. Variable selection using random forests. Pattern Recognition Letters, 2010.

11. Da Wei Huang, Brad T Sherman, Richard A Lempicki, et al. Systematic and integrative analysis of large gene lists using david bioinformatics resources. Nature protocols, 4(1):44–57, 2008.

12. H Ishwaran, U B Kogalur, and E H Blackstone. Random survival forests. The Annals of Applied Statistics (2008): 841-860., 2008.

13. SY Jung, Junsoo Jeong, Seung-Ho Shin, Youngmee Kwon, Eun-A Kim, Kyoung Lan Ko, Kyung Hwan Shin, Keun Seok Lee, In Hae Park, Seeyoun Lee, Seok Won Kim, Han-Sung Kang, and Jungsil Ro. The invasive lobular carcinoma as a prototype luminal A breast cancer: A retrospective cohort study. BMC Cancer, 10(1):664, December 2010.

14. Brian D Lehmann, Joshua A Bauer, Xi Chen, Melinda E Sanders, A Bapsi Chakravarthy, Yu Shyr, and Jennifer A Pietenpol. Identification of human triple-negative breast cancer subtypes and preclinical models for selection of targeted therapies. The Journal of clinical investigation, 121(7):2750–2767, July 2011.

15. Stefan Michiels, Serge Koscielny, and Catherine Hill. Prediction of cancer outcome with microarrays: a multiple random validation strategy. The Lancet, 365(9458):488–492, February 2005.

16. Vamsi K Mootha, Cecilia M Lindgren, Karl-Fredrik Eriksson, Aravind Subramanian, Smita Sihag, Joseph Lehar, Pere Puigserver, Emma Carlsson, Martin Ridderstråle, Esa Laurila, Nicholas Houstis, Mark J Daly, Nick Patterson, Jill P Mesirov, Todd R Golub, Pablo Tamayo, Bruce Spiegelman, Eric S Lander, Joel N Hirschhorn, David Altshuler, and Leif C Groop. PGC-1alpha-responsive genes involved in oxidative phosphorylation are coordinately downregulated in human diabetes. Nature genetics, 34(3):267–273, July 2003.

17. The Cancer Genome Atlas Network. Comprehensive molecular characterization of human colon and rectal cancer. Nature, 487(7407):330–337, July 2012.

18. Jennifer Pittman, Erich Huang, Holly Dressman, Cheng-Fang Horng, Skye H Cheng, Mei-Hua Tsou, Chii-Ming Chen, Andrea Bild, Edwin S Iversen, Andrew T Huang, Joseph R Nevins, and Mike West. Integrated modeling of clinical and gene expression information for personalized prediction of disease outcomes. PNAS, 101(22):8431–8436, June 2004.

19. Yanjun Qi, Ziv Bar-Joseph, and Judith Klein-Seetharaman. Evaluation of different biological data and computational classification methods for use in protein interaction prediction. Proteins: Structure, Function, and Bioinformatics, 63(3):490–500, 2006.

20. G Riddick, H Song, S Ahn, J Walling, D Borges-Rivera, W Zhang, and H A Fine. Predicting in vitro drug sensitivity using Random Forests. Bioinformatics, 27(2):220–224, January 2011.

21. Xavier Sastre Garau, Michel Jouve, Bernard Asselain, Anne Vincent Salomon, Philippe Beuzeboc, Thierry Dorval, Jean Claude Durand, Alain Fourquet, and Pierre Pouillart. Infiltrating lobular carcinoma of the breast: Clinicopathologic analysis of 975 cases with reference to data on conservative therapy and metastatic patterns. Cancer, 77(1):113–120, January 1996.

22. Kenjiro Sawada, A Reza Radjabi, Nariyoshi Shinomiya, Emily Kistner, Hilary Kenny, Amy R Becker, Muge A Turkyilmaz, Ravi Salgia, S Diane Yamada, George F Vande Woude, Maria S Tretiakova, and Ernst Lengyel. c-Met overexpression is a prognostic factor in ovarian cancer and an effective target for inhibition of peritoneal dissemination and invasion. Cancer research, 67(4):1670–1679, February 2007.

23. Kristy Stengel and Yi Zheng. Cdc42 in oncogenic transformation, invasion, and tumorigenesis. Cellular Signalling, 23(9):1415–1423, September 2011.

24. Aravind Subramanian, Pablo Tamayo, Vamsi K Mootha, Sayan Mukherjee, Benjamin L Ebert, Michael A Gillette, Amanda Paulovich, Scott L Pomeroy, Todd R Golub, Eric S Lander, and Jill P Mesirov. Gene set enrichment analysis: a knowledge-based approach for interpreting genome-wide expression profiles. PNAS, 102(43):15545–15550, October 2005.

25. Kimita Suyama, Irina Shapiro, Mitchell Guttman, and Rachel B Hazan. A signaling pathway leading to metastasis is controlled by N-cadherin and the FGF receptor. Cancer cell, 2(4):301–314, October 2002.

26. Robert Tibshirani. Regression shrinkage and selection via the lasso. Journal of the Royal Statistical Society. Series B (Methodological), pages 267–288, 1996.

27. Wouter G Touw, Jumamurat R Bayjanov, Lex Overmars, Lennart Backus, Jos Boekhorst, Michiel Wels, and Sacha A F T van Hijum. Data mining in the Life Sciences with Random Forest: a walk in the park or lost in the jungle? Briefings in bioinformatics, 14(3):315–326, May 2013.

28. Gulisa Turashvili, Jan Bouchal, George Burkadze, and Zdenek Kolar. Differentiation of tumours of ductal and lobular origin: II. Genomics of invasive ductal and lobular breast carcinomas. Biomedical papers of the Medical Faculty of the University Palacký, Olomouc, Czechoslovakia, 149(1):63–68, June 2005.

29. A Villanueva, Y Hoshida, C Battiston, V Tovar, and D Sia. Combining clinical, pathology, and gene expression data to predict recurrence of hepatocellular carcinoma. Gastroenterology, 2011.

30. Ming Yuan and Yi Lin. Model selection and estimation in regression with grouped variables. Journal of the Royal Statistical Society: Series B (Statistical Methodology), 68(1):49–67, 2006.

